# DSPE-PEG does not retain targeting antibodies on LNP surfaces *in vivo*; a higher molecular weight anchor is required

**DOI:** 10.64898/2026.07.02.736109

**Authors:** Brian K. Wilson, Lucas D. Johnson, Jason Liu, Nicholas Caggiano, Nikhil Subraveti, Karthik Nagapudi, Andrew Tsourkas, Robert K. Prud’homme, Kurt D. Ristroph

**Author notes:** equal contribution (Brian K. Wilson, Lucas D. Johnson).

## Abstract

Extrahepatic delivery of lipid nanoparticles (LNPs) to non-phagocytic cells is a major challenge, with the leading strategy involving surface functionalization with target-specific monoclonal antibody (mAb) ligands. We investigate the stability of mAb-conjugated LNPs using two anchoring systems: the commonly used DSPE-PEG_2kDa_-maleimide and a block copolymer, PCL_5kDa_-b-PEG_2kDa_ -maleimide, with the hypothesis that conjugation to a 150,000 Da antibody could overwhelm the relatively small ∼600 Da aliphatic anchor on the PEG-lipid *in vivo*. Shedding of the mAB would compromise targeting. Conjugation integrity following IV injection was assessed by tagging LNPs and mAbs with metal ion tracers that could be quantified by ICP-MS. Results show that DSPE-PEG-mAb rapidly (within 1h) dissociates from LNPs in blood, leading to accelerated LNP clearance. In contrast, mAbs conjugated using PCL-b-PEG remained stably associated with the LNP over the 24h circulation and clearance of the construct. Results are connected to a thermodynamic model that reproduces experimental findings for PEG-anchor(-mAb) shedding *in vitro* and *in vivo*. This study identifies anchoring strength as a critical, unconsidered parameter for *in vivo* performance when conjugating mAbs to LNPs for extrahepatic delivery.

**Graphical abstract:** 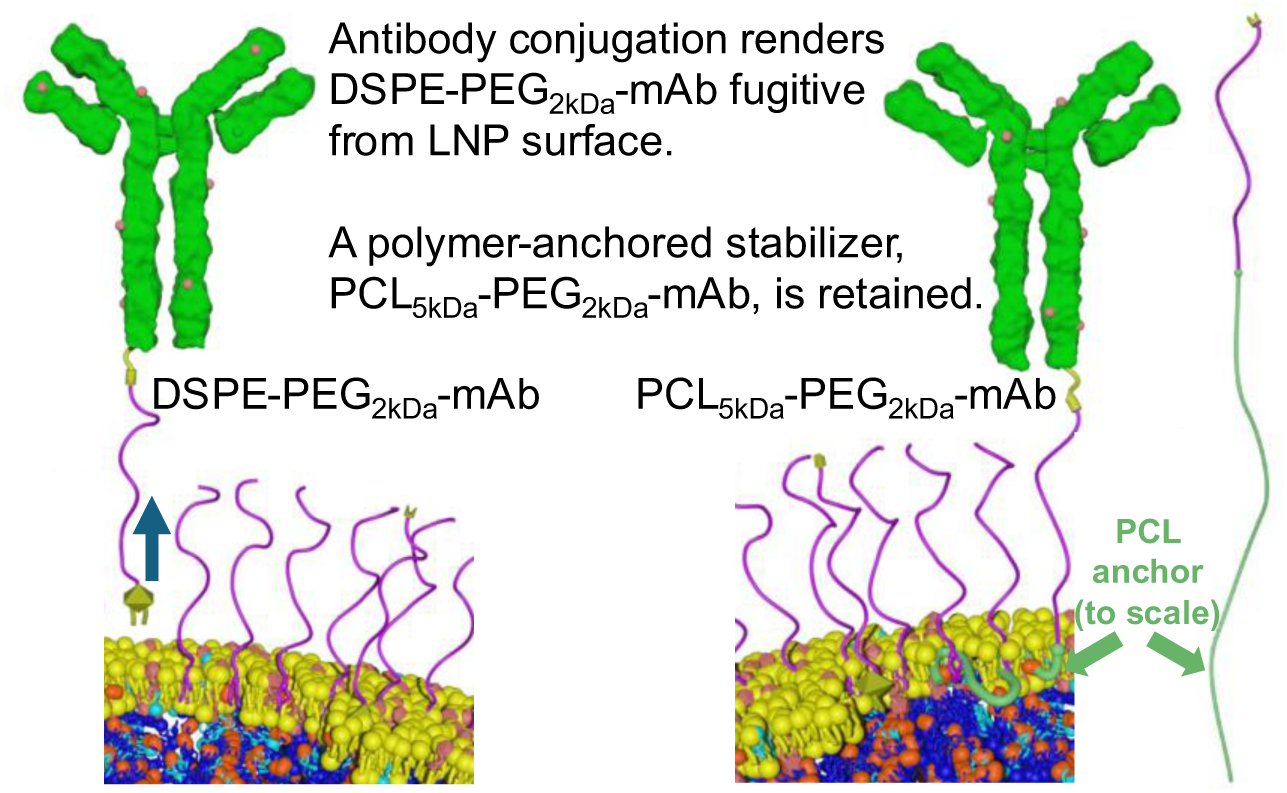

## Body

Nucleic acids are potent therapeutic molecules with immense therapeutic potential but require a sophisticated intracellular delivery strategy. Lipid nanoparticles (LNPs) are the most well-developed class of nucleic acid delivery vehicle, having demonstrated intracellular delivery of payloads ranging from low molecular weight antisense oligonucleotides to extremely high molecular weight DNA plasmids.^1^ Clinical use of LNPs has expanded significantly in the last 5 years, notably in combating infectious disease threats through successful induction of neutralizing antibody responses to immunogenic epitopes.^2^

Existing commercial LNPs contain 4 lipid components: an ionizable lipid, a zwitterionic structural lipid, cholesterol, and a poly(ethylene glycol)-lipid (PEG-lipid) stabilizer. Each lipid component plays a key role in the physical structure of the LNP, which evolves significantly between initial self-assembly in an acidic pH buffer and subsequent processing into a neutral pH buffer (**Figure 1a**). Ionizable lipids are cationic in the acidic assembly media (typically pH ∼4), which promotes nucleic acid complexation and produces small, vesicle-like cationic LNPs with both excess charged ionizable lipid and PEG-lipid together electrosterically stabilizing the large specific surface area.^3^ Exchanging the acidic buffer for a neutral buffer (pH 7.4) causes the majority of the ionizable lipid to revert to the uncharged free base form, leaving only the sparse PEG-lipid as the stabilizer on the LNP surface.^4,5,6^ This PEG density is insufficient to stop particle-particle coalescence, so LNP fusion occurs until the PEG density recovers sufficiently to prevent further LNP-LNP fusion (**Figure 1a**). As a result, LNPs transform from small, electrosterically-stabilized vesicle-like particles at acidic pH into larger, sterically-stabilized LNPs with an oil-like core mainly consisting of uncharged ionizable lipid at neutral pH (**Figure 1a**).^7^

**Figure 1:**
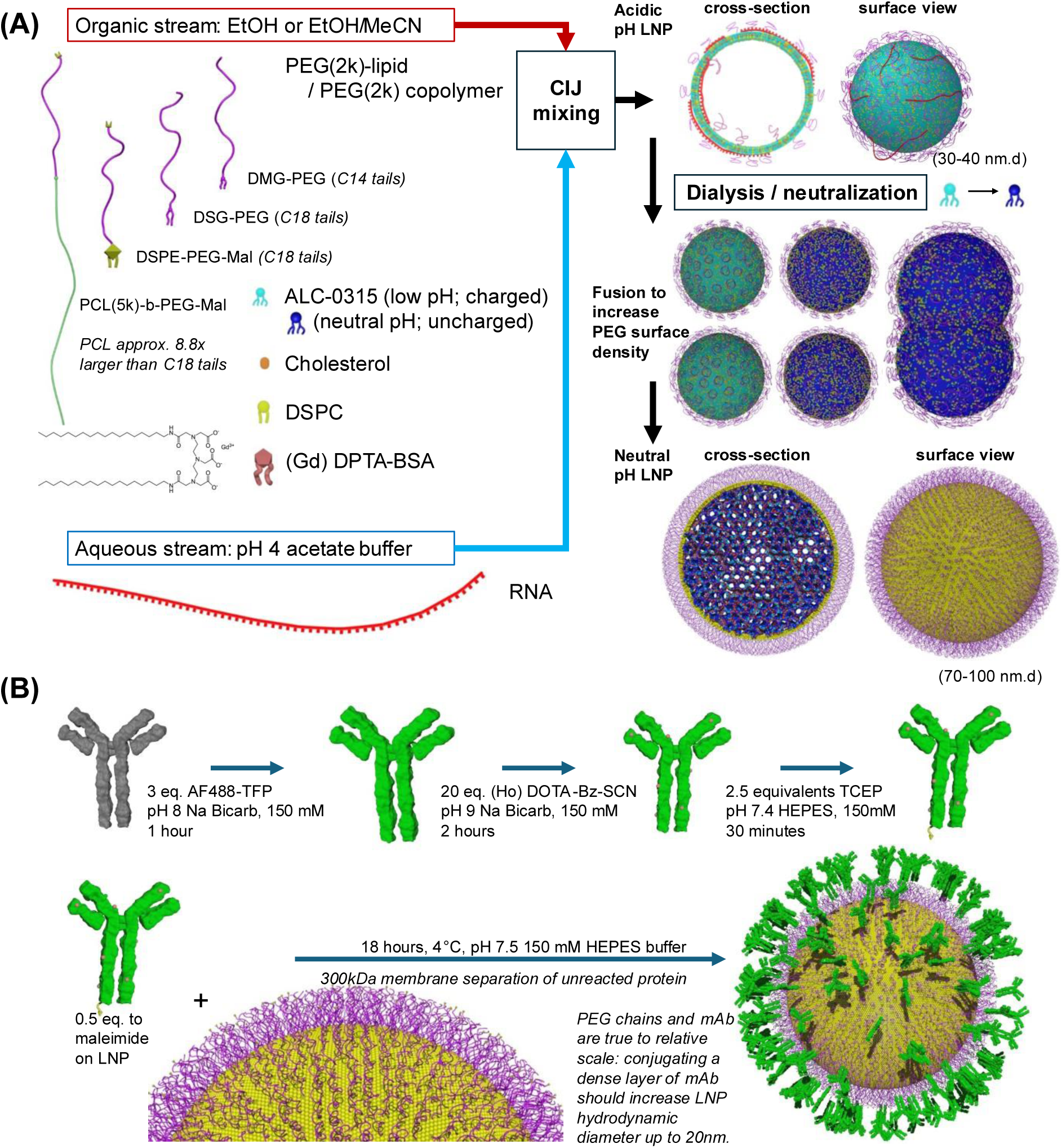
LNP formulation, processing, structure, and reaction scheme. (A) LNPs are formed by antisolvent precipitation using a scalable turbulent-flow mixer and then neutralized over dialysis, followed by (B) antibody conjugation.

The PEG-lipid stabilizer ultimately controls LNP size and stability, critical quality attributes for particle-based delivery systems, through the eventual formation of a sufficiently dense PEG layer on the LNP surface during formation and processing. *In vivo*, the clinically-used PEG-lipid stabilizer, dimyristoyl-glycero-PEG_2kDa_ (DMG-PEG), rapidly partitions off LNP surfaces when exposed to complex biological fluids due to the low adsorption energy of its short twin 14-carbon alkyl tails.^8^ Shedding of the PEG-lipid stabilizer and subsequent formation of a protein corona is advantageous in immunogenic applications, where the adsorbed proteins promote LNP trafficking to and protein translation in phagocytotic cells, a critical early step in eliciting antibody responses against the antigen encoded by the nucleic acid payload.^9^ This intentional use of a fugitive PEG-lipid stabilizer means current commercial LNP formulations struggle to avoid uptake and clearance from circulation by phagocytotic or peripheral blood mononuclear cells. PEG layers anchored with longer alkyl tails, such as the twin saturated 18-carbon long tails of distearoyl glycerol-PEG_2kDa_ (DSG-PEG), are not fugitive from the LNP surface, producing a vehicle with improved stability and tripling of blood circulation half-life – though LNPs with these strongly-anchored PEG-lipid surfaces still tend to eventually accumulate in the liver and spleen.^8^

Conjugation of an antibody to the PEG chains on an LNP surface through the DSPE-PEG lipid is a common strategy proposed to prepare targeted LNPs.^10,11,12,13,14,15^ Antibody conjugation requires adding to the formulation a reactive PEG-lipid as a fifth component, typically in the form of a distearoyl-glycero-phosphatidyl ethanolamine-PEG (DSPE-PEG) derivative with a reactive end group, such as DSPE-PEG-maleimide. This reactive-PEG-bearing LNP is then conjugated to a suitably derivatized mAb (e.g. a thiolated mAb for DSPE-PEG-maleimide). Reaction conditions must be optimized to preserve the size and polydispersity of the initial LNPs, and oftentimes mAb conjugation reactions yield LNP populations with aggregated, polydisperse particle size distributions. Our experience with using DSPE-PEG-maleimide to anchor thiolated mAbs onto LNPs is that the reaction requires a high salinity environment to avoid LNP aggregation. As soon as the ionic strength is decreased, or hydrophobic “sink” conditions exist, the DSPE-PEG-mAb LNPs become unstable and aggregate into much larger particles, rather than the ∼20nm increase in size that would be expected from addition of a mAb to the LNP surface. This instability only occurs after conjugating mAb to the DSPE-PEG, leading to our present hypothesis for this work: that conjugating a large mAb (about 150,000 Da) to a small DSPE-PEG (about 2,700 Da, less than 600 Da of which is in the anchoring hydrophobic tails) overwhelms the anchoring potential of the DPSE 18-carbon alkyl tails, rendering the DSPE-PEG-mAb fugitive from the particle surface (analogous to the 2,000 Da PEG chain in DMG-PEG overwhelming the two 14-carbon tails and rendering that molecule labile from the surface) when exposed to *in vivo* environments., This in turn would substantially reduce the time window for targeting, thereby rendering this strategy for extrahepatic delivery ineffective.

Amphiphilic block copolymers present a natural solution to the problem of strongly anchoring a large mAb to an LNP surface. Poly(ester)-block-poly(ethylene glycol) polymers are biocompatible, degradable materials commonly used in the production of biomedical nanoparticles.^16^ Poly(caprolactone)-block-poly(ethylene glycol) (PCL-b-PEG) has desirable solubility and miscibility behavior with the lipids used in conventional 4-component formulations and can be substituted into the LNP formulation without compromising LNP colloidal properties or nucleic acid polymer delivery.^17^

In this study, we present the preparation of cetuximab-conjugated LNPs and measurement of their circulating time after intravenous injection into Sprague-Dawley rats. Whereas Mui, Cullis *et al.* used radioactive labeling to track the *in vivo* fate of the lipids^8^, here we demonstrate the use of inductively coupled plasma mass spectrometry (ICP-MS) tracking using lanthanide tracers. Materials chelating different lanthanides were added to the LNP vehicle and onto the mAb, to make each component detectable by ICP-MS (**Figure 1**). In addition to the usual 4-component lipids (ALC-0315 ionizable lipid, DSPC zwitterionic lipid, cholesterol, and DSG-PEG stabilizer) our LNPs also contain a gadolinium-chelating DTPA-bistearylamide (Gd-BSA)^18^ lipid and one of either DSPE-PEG_2kDa_-maleimide or PCL_5kDa_-b-PEG_2kDa_-maleimide as the reactive component to conjugate thiolated cetuximab. As an aside regarding the use of DSG-PEG: at 1.5%mole, using non-fugitive DSG-PEG as the surface stabilizer instead of the fugitive DMG-PEG has been shown not to interfere with LNP biological activity; only at higher mole fractions does the higher molecular weight anchor compromise activity, likely by interfering with endosomal escape^8^. It has also been shown that even 1.5%mole PCL-b-PEG does not compromise LNP bioactivity^17^. The cetuximab was derivatized via its amines with holmium-loaded DOTA-benzyl-isothiocyanate (Ho-DOTA) (**Figure 1b**).^19^ Under this scheme, both the LNP core, with the Gd-BSA, and the attached mAb, with the Ho-DOTA, can be quantitated in blood and tissue samples using ICP-MS. Measurements of Gd and Ho in blood over time show that for LNPs with DSPE-PEG-mAb, the LNP-associated Gd signal rapidly clears into the liver and spleen, while the mAb-associated Ho signal circulates in blood as it is slowly cleared over 24 hours. For LNPs with PCL-b-PEG-mAb, mAb-associated Ho and LNP-associated Gd signals remain together over 24 hours as the LNPs are slowly cleared from circulation. This is consistent with the hypothesis of DSPE-PEG-mAb partitioning from the LNP surface and demonstrates that block copolymers present a route for the development of robust, antibody-conjugated LNPs.

LNPs used in this study are variants on the usual Patisiran (Onpattro®) formulation commonly encountered as a ‘standard’ LNP but with an additional gadolinium-chelating lipid replacing a fraction of the lipid components. The base formulation is 50% mole ALC-0315, 37.5% mole cholesterol, 10% mole DSPC, 1.5% mole PEG-lipid stabilizer, and 2% mole gadolinium-EDTA-bistearylamide. The specific formulations have different compositions of the PEG-lipid stabilizer, with a reactive PEG-maleimide-bearing stabilizer added for protein conjugation. Briefly, the four formulations have stabilizer compositions of: [1] 1.2% mole DSG-PEG plus 0.3% mole DSPE-PEG-maleimide (“DSPE-PEG-mAb”); [2] 1.2% mole DSG-PEG plus 0.3% mole PCL-b-PEG-maleimide (“PCL-b-PEG-mAb”); [3] 1.5% mole DSG-PEG (“DSG”); or [4] 1.5% mole PCL-b-PEG (“PCL”). The last two formulations are untargeted controls included to demonstrate the impact of the DSG versus PCL PEG-anchoring groups on LNP circulation times (**Table 1**).

**Table 1:** LNP formulations, size/PDI, and metal loading.

|  | F-1, DSPE-PEG-mAb | F-2, PCL-b-PEG-mAb | F-3, DSG-PEG (control) | F-4, PCL-b-PEG (control) |
| --- | --- | --- | --- | --- |
| (all in mol%) |  |  |  |  |
| ALC-0315 | 50 | 50 | 50 | 50 |
| DSPC | 10 | 10 | 10 | 10 |
| Chol. | 35.9 | 35.9 | 35.9 | 35.9 |
| (Gd)DTPA-BSA | 2 | 2 | 2 | 2 |
| EtTP-5 | 0.1 | 0.1 | 0.1 | 0.1 |
| DSG-PEG | 1.2 | 1.2 | 1.5 | 0 |
| PCL-b-PEG | 0 | 0 | 0 | 1.5 |
| DSPE-PEG-Mal | 0.3 | 0 | 0 | 0 |
| PCL-b-PEG-Mal | 0 | 0.3 | 0 | 0 |
| Diameter, nm (post-conjugation) | 69.5 ± 1.4<br>(87.6 ± 1.7) | 76.6 ± 0.8<br>(92.7 ± 1.6) | 68.1 ± 1.0<br>(NA) | 96.3 ± 0.9<br>(NA) |
| PDI (post-conjugation) | 0.158<br>(0.017) | 0.100<br>(0.013) | 0.114<br>(0.008) | 0.123<br>(0.012) |
| ng Gd/mL | 19895.95 | 26694.35 | 25998.05 | 21867.2 |
| ng Ho/mL | 39752.8 | 42063.55 | 0.15 | 0.4 |
| Mass ratio, Gd:Ho | 0.50 | 0.63 | N/A | N/A |

All LNPs were synthesized by Flash NanoPrecipitation (FNP) in a confined impinging jets mixer (CIJ) from a 6 mg/mL solution of lipids (70/30 v/v EtOH/THF) mixed against a 20 mM pH 4 acetate buffer containing poly(adenylic acid) (polyA) as the model nucleotide for encapsulation at an N:P ratio of 6.^20^ Mixer effluent was directed into a quench bath containing 10 mM acetate buffer at pH 4 with a total volume 8 times that of the ethanolic lipid stream; for example, 20 mL of a 6 mg/mL ethanol stream of lipids was mixed against 20 mL of acetate buffer antisolvent containing polyA with collection in a 160 mL bath of acetate buffer.^21^ The quenched LNPs were then concentrated via centrifugal ultrafiltration (Amicon filter, 100 kDa MWCO, 500 RCF, approx. 90 minutes total; filters were first rinsed with 15mL of 30/70 EtOH/water v/v at 500rcf followed by 15mL of deionized water twice) to reduce the total volume from 200 mL to approximately 15 mL. This 15 mL concentrated LNP dispersion was then loaded into 30 kDa MWCO dialysis cassettes and first dialyzed against 1000 mL of 10 mM HEPES at pH 7.4 containing 10% volume ethanol, then in a second step dialyzed against 1000 mL of 10 mM HEPES at pH 7.4, each step taking two hours. This two-step process was necessary for preserving the small average size and polydispersity of LNPs containing DSPE-PEG-maleimide. LNP hydrodynamic sizes were measured by dynamic light scattering (DLS).

All LNP formulations exhibited similar average diameters around 70-80 nanometers post dialysis, as shown in **Figure 2**. The control group with only PCL-b-PEG stabilizer yielded slightly larger LNPs (∼95nm) after the dialysis workup, which is consistent with the larger PCL anchoring block footprint lowering the PEG surface density compared to the smaller DSG anchor, which allows LNPs to grow larger during the neutralization-mediated fusion process in dialysis. Average sizes and properties of each LNP formulation are given in **Table 1**.

**Figure 2:**
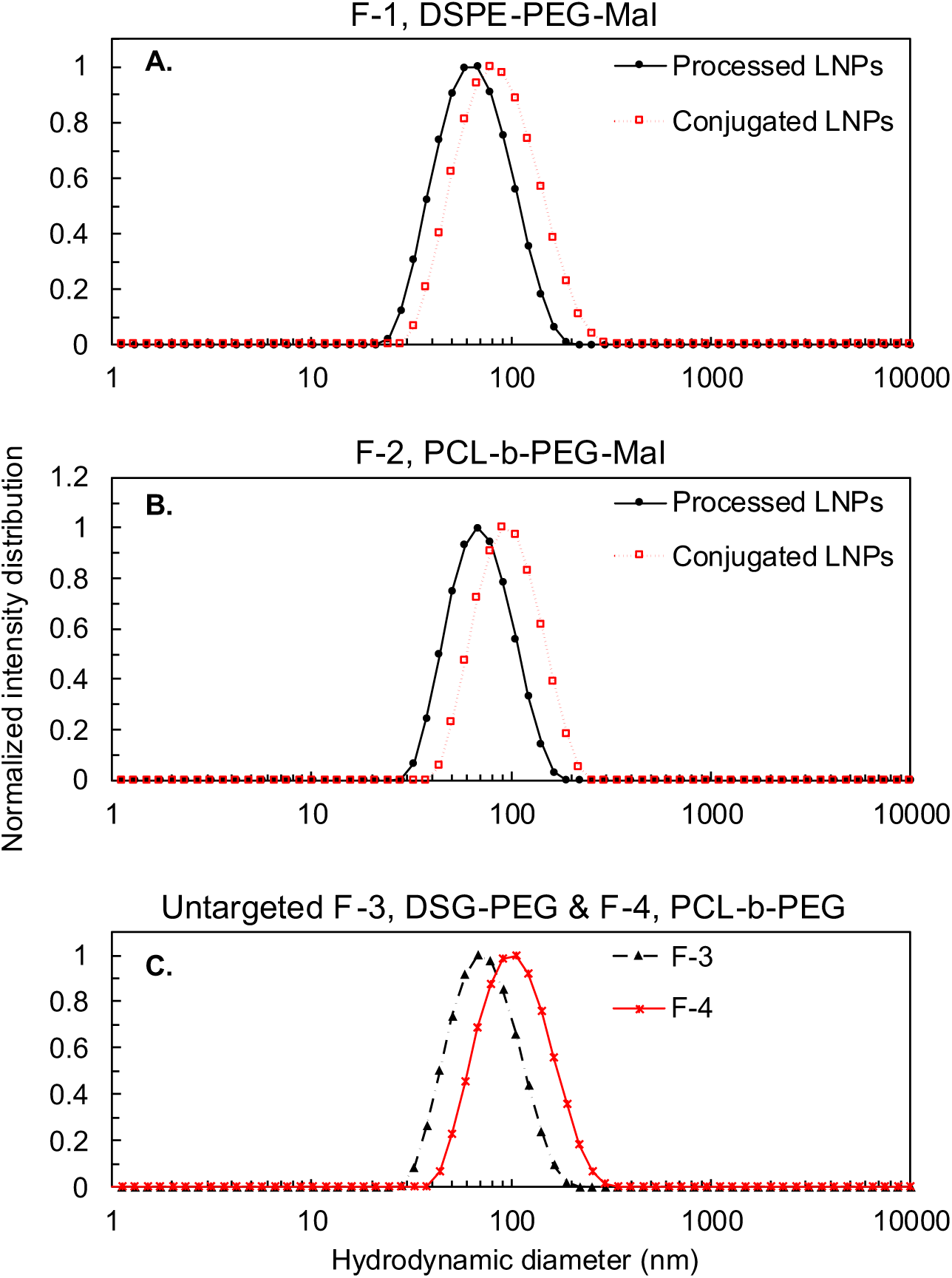
Production of large-scale LNP batches and conjugation to monoclonal antibody demonstrated for 50%mol ALC-0315, 35.9% cholesterol, 10% DSPC, and 2% (Gd)-DTPA-BSA formulations containing 1.2% DSG-PEG_2kDa_ and 0.3% either DSPE-PEG_2kDa_-maleimide or PCL_5kDa_-b-PEG_2kDa_-maleimide conjugated to thiolated holmium-tagged antibodies. LNPs were made in 250 mL batches, concentrated to approximately 20 mL, and dialyzed against neutral buffer then characterized. (A) Particle size distribution for LNPs with 0.3%mol DSPE-PEG_2kDa_-maleimide before (solid circles) and after (open squares) conjugation. (B) Particle size distribution for LNPs with 0.3%mol PCL_5kDa_-b-PEG_2kDa_-maleimide before (solid circles) and after (open squares) conjugation.

The two-step dialysis process (change pH at 10% EtOH, then remove EtOH at constant pH 7.4) was necessary to avoid significant growth of the 0.3% mole DSPE-PEG-maleimide formulation. This formulation exhibited the same small initial size (approximately 45 nm diameter) at pH 4 as the other formulations, but attempting to directly dialyze – i.e. remove ethanol and change pH simultaneously – caused aggregation to a diameter above 150 nm (**SI Figure S1**). DSPE-PEG is anionic, whereas the DSG-PEG and PCL-b-PEG stabilizers are neutral. DSPE begins as a zwitterionic phosphatidyl (anion) ethanolamine (cation) lipid, but the PEG chain is conjugated as an amide onto the terminal ethanolamine. This leaves only the negative charge of the phosphatidyl group, making DSPE-PEG an anionic surfactant. The combination of intrinsic curvature of this anionic surfactant, the large excess of cationic ionizable lipids, and the need for primary LNPs to undergo rearrangement during the neutralization-fusion process means that DSPE-PEG destabilizes the LNP during dialysis and leads to particle growth by aggregation.^22^ Performing the pH-mediated fusion process in the presence of 10% EtOH preserved the LNP size by the alcohol cosolvent increasing the fluidity of the LNP surface.

Cetuximab (as Erbitux® solution) was modified with a covalently conjugated Holmium chelator to render the protein detectable by ICP-MS. Additional details regarding the conjugation can be found in the Supplementary Information.

TCEP-activated cetuximab-Ho was mixed with maleimide bearing LNPs at a molar ratio of 0.5:1 mAb:maleimide. After 12 hours, the reaction mixture was placed in a 300 kDa MWCO float-a-lyzer and dialyzed against 10 mM HEPES pH 7.4, 140 mM NaCl for 18 hours. LNP size was remeasured and showed an approximate 20 nanometer increase over the parent unconjugated LNPs, suggesting a significant extent of mAb conjugation to these formulations (**Figure 2** and **Table 1**).

Buffer conditions used for protein conjugation reactions play a critical role in determining the stability of the adduct on the LNP surface. Performing the DSPE-PEG-maleimide conjugation at low salt conditions (10 mM HEPES buffer only) led to the rapid formation of aggregates and a large increase in particle size measured by DLS. Increasing the total ionic strength to 150 mM (10 mM HEPES, 140 mM NaCl) arrested this aggregation and led to a controlled, roughly 20 nm increase in particle size, as expected from the addition of a mAb layer on the LNP surface. The higher ionic strength screens the electrostatic interactions between the charged DSPE-PEG and decreases its solubility in the buffer. In contrast, the neutral PCL-b-PEG-maleimide system showed no sensitivity to the total salinity (e.g. 10mM versus 150mM), with either condition yielding stable LNPs that were about 15-20 nanometers larger post-conjugation.

LNPs were administered via tail vein injection to male Sprague-Dawey rats (n=5 per treatment) at 7.5mg LNP/kg. Serial blood draws at 1, 3, 7, 10, and 24 hours were collected for analysis of the circulating particle quantity and composition by ICP-MS. After the 24-hour draw, animals were sacrificed, and the brain, heart, lungs, spleen, and liver were collected for analysis. Animal tissue and whole-blood samples were digested using a 3:1:1 (v/v) mixture of 70% nitric acid, hydrogen peroxide, and hydrochloric acid in 75 mL Teflon MARSXpress vessels and subjected to microwave-assisted digestion in the MARS 6 Microwave Digestion System following the One Touch method for “Wet Animal Tissue”: a single-step ramp to 200°C in 15 minutes, and hold for 15 minutes at 800 psi and 900-1800W. For every digestion batch, three reagent-only blank vessels and three reference samples (NIST 1577 C “Bovine Liver”) were included. All samples received an offline addition of Europium, with a final concentration of 10 µg/L Eu after dilution. Following digestion, samples were diluted to 18% nitric acid, which was clear and particle-free, and filtered through 1.5 µm nylon membrane filters. Total metal content was quantified using ICP-MS (Agilent 7850). Additional details regarding the cleaning of the digestion vessels and ICP-MS parameters can be found in the Supplementary Information.

Measured quantities of Holmium (mAb) and Gadolinium (LNP), converted to percentage of the initial dose (%ID), in the serial blood draws at each time point are shown in **Figure 3**. The mAb-conjugated LNPs show detectable Ho and Gd signals, while the control DSG-PEG and PCL-b-PEG groups (without mAb) show only measurable Gd signals. Data from the mAb-conjugated formulations is also plotted as the mass ratio of Gd:Ho to better visualize changes in the circulating material over time (**Figure 3e-f**). Each formulation was characterized for its total metal content before administration (**Table 1**), which provides a characteristic Gd:Ho ratio for the intact formulation.

**Figure 3:**
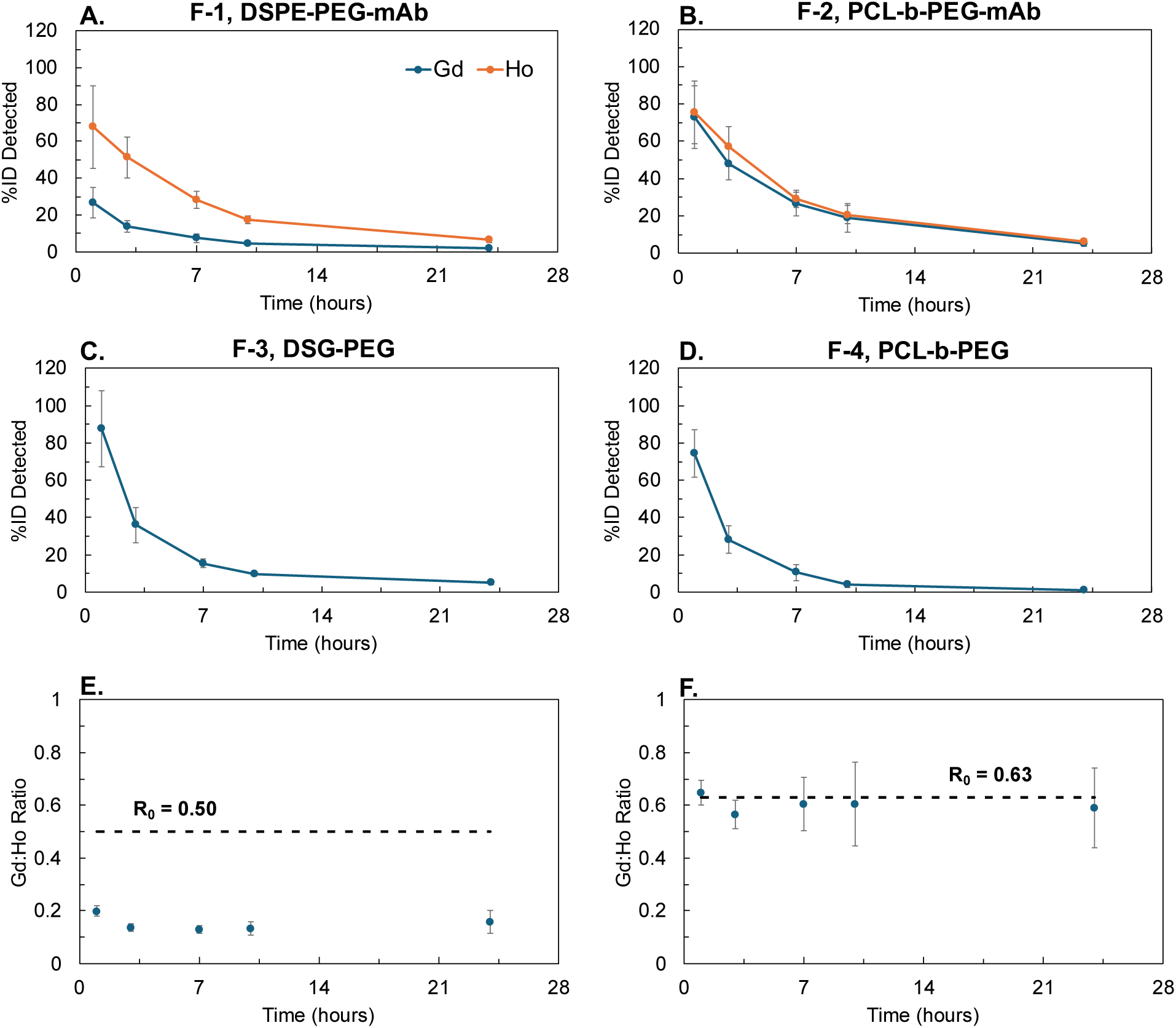
(A-D) Concentrations of Ho and Gd detected in whole blood over time. (E-F) Gd-Ho ratio at each timepoint of panels A (E) and B (F). Changes in the Gd:Ho ratio suggest dissociation and clearance of one component of the mAb-conjugated LNP formulation. The DSPE-PEG-mAb shows a change from its initial Gd:Ho ratio (Table 1: 0.50 for formulation F1, 0.63 for formulation F2) by the first time point at 1h, while the PCL-b-PEG-cetuximab shows a constant Gd:Ho ratio over 24 hours. The change in ratio for the mAb-PEG-DSPE LNP is due to dissociation of the DSPE-PEG-cetuximab and subsequent clearance of the Gd-bearing LNP portion, as evidenced by the sustained Ho circulation over 24 hours. In contrast, the PCL-b-PEG-mAb LNPs remain intact, with the tagged mAb and tagged LNP clearing at the same rate.

There are three significant conclusions to draw from the four sets of blood draw data:

First, the block copolymer-anchored mAb remains attached to the NP, with no indication of shedding during the entire 24 hour duration of the test. The ratio of Gd:Ho remains constant at the initial value (0.63) during the 24 hours of circulation (**Figure 3b**). The PCL-b-PEG-mAb LNP clears relatively slowly and remains intact over 24 hours, which satisfies the circulation time deemed necessary in the liposomal delivery field to reach hard-to-engage tumors or small metastases.

Second, for the DSPE-anchored mAb: by the first time point of 1 hr, only 27% of the LNP core is still in the blood volume (**Figure 3a**). It has been cleared, predominantly to the liver (**Figure 4**). When the mAb partitions off the LNP surface, it removes 20% of the surface PEG with it, such that proteins such as ApoE quickly adsorb and direct clearance to the liver. The DSPE-PEG-mAb shows a change from its initial Gd:Ho ratio of 0.50 at injection to 0.2 at 1 hr (**Figure 3e**). This change in ratio is due to dissociation of the DSPE-PEG-mAb and subsequent clearance of the Gd-bearing LNP core. The continued high level of tagged mAb in the blood indicates that the partitioned mAb has associated with lipoproteins, RBC membranes, or other hydrophobic circulating sinks. This result contradicts the currently held belief that PSPE-PEG-mAb will remain anchored in the LNP during extended circulation times.

**Figure 4:**
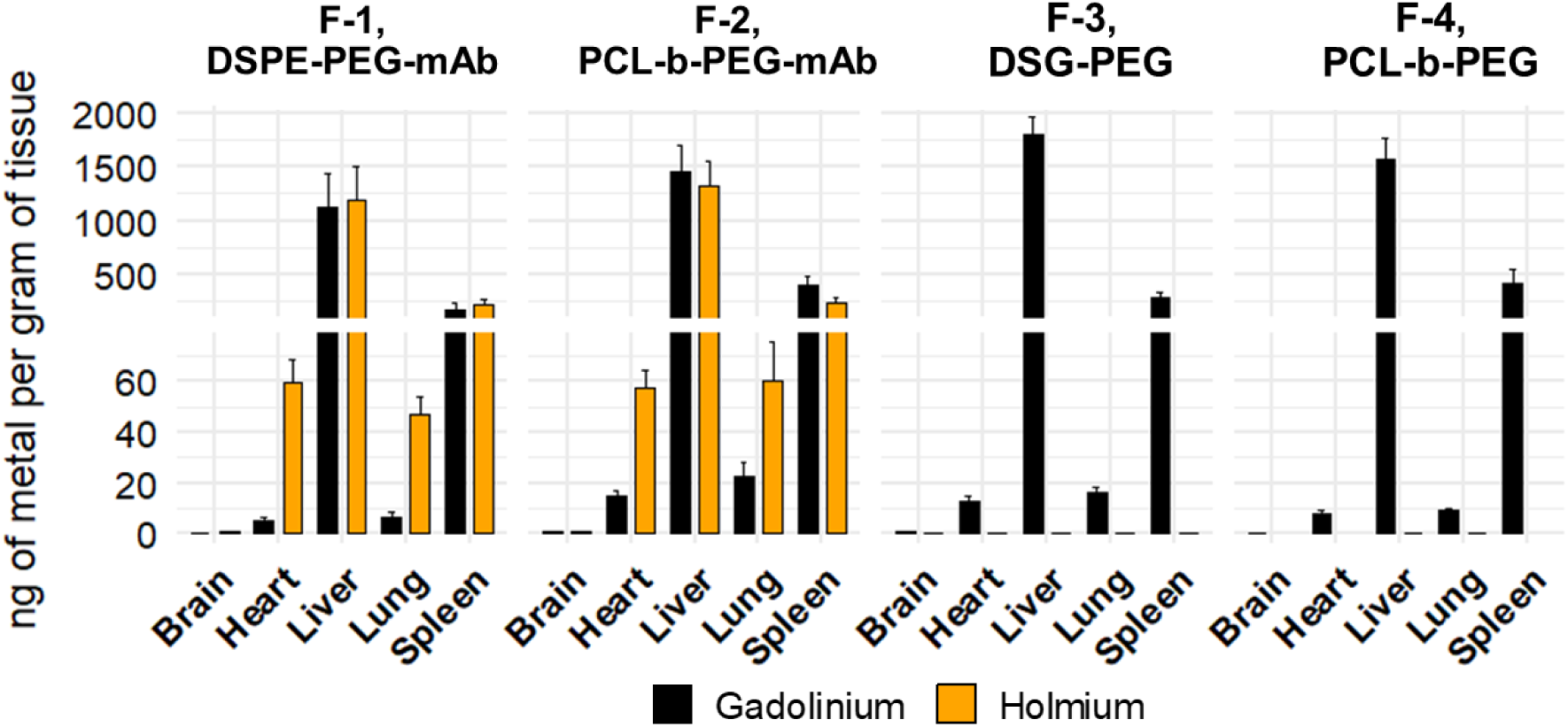
Metal distributions in organs after final time point. No major differences are observed across the formulations.

Third, for the two controls, LNPs stabilized with DSG-PEG and PCL-b-PEG, the PEG layer is stable. The circulation half life for both is ∼2 hrs, shorter than what has been achieved by “stealth” nanoparticles with denser surface PEG layers, which have achieved circulation times on the order of 24 hrs.^23,24^ The shorter circulation time is because the PEG conformation on LNPs with these or similar compositions has been reported as around 0.273 chains/nm^2^, near the upper end of the ‘mushroom’ regime but below the density required for a truly densely-packed PEG ‘brush’ required to prevent protein adsorption and for extended circulation^25^, but which may interfere with endosomal escape if installed onto LNPs.^5,8^

The fate of the Gd-tagged LP and Ho-tagged mAb was determined after the final 24-hour blood draw time point by harvesting and analyzing the major organs. As expected, most of the LNP-associated signal ends in the liver and spleen (**Figure 4, SI Table S3**), as the study was done using healthy rats with no tumor to target. There were no major statistically significant differences in organ accumulation between the different LNP formulations.

PEG-lipids serve as the anchors attaching antibodies to the surface of a targeted LNP. Bio-orthogonal conjugation chemistries attach the mAb, often nonspecifically as done here, to the PEG-lipid to form a large, lipid-PEG-mAb construct. Addition of the mAb radically changes the molecular weight of the PEG-lipid by attaching a 150,000 Da biomacromolecule to a 2,000 – 10,000 Da PEG-lipid molecule. Antibodies have colloidal properties due to their large molecular weights and globular structure; an antibody is a roughly 11 nanometer diameter nanoparticle composed of four disulfide crosslinked protein polymer chains. Bringing a mAb into proximity with the LNP surface reduces the translational entropy of the mAb and the PEG tether. This creates a free energy penalty for remaining surface-bound, and, at high surface coverage, excluded volume effects between mAbs introduce an additional entropic penalty. A stable anchor-PEG-mAb conjugate requires sufficiently strong anchoring to offset these entropic losses and prevent desorption.

DSPE-PEG-maleimide and PCL-b-PEG-maleimide exhibited completely different stability behaviors during the conjugation process to thiolated proteins, either bovine serum albumin (BSA) or cetuximab. A low (10mM, HEPES buffer ions only) ionic strength pH 7.4 media caused rapid aggregation of the DSPE-PEG-protein-stabilized LNPs via the PEG-lipid conjugate pulling out of the LNP surface (**SI Figure S2**). A high (150 mM total, 10 mM HEPES buffer ions and 140 mM NaCl) ionic strength media produced stable (to *in vitro* conditions) DSPE-PEG-protein bearing LNPs. Both intermediate (65,000 Da BSA) and large (150,000 Da cetuximab) protein conjugates showed this same behavior in which high ionic strength was required to avoid LNP aggregation after the protein conjugation reaction with DSPE-PEG-maleimide. In contrast, the PCL-b-PEG-maleimide system resulted in stable, protein-modified LNP surfaces in either high or low ionic strength media. Increased salt concentration makes withdrawing the hydrophobic lipid tail and charged phosphate residue into the aqueous environment less favorable (seen experimentally by a higher critical micelle concentration (CMC) for DSPE-PEG in water than in buffer^26^) and promotes retention of the DSPE-PEG-protein conjugate, while the PCL-b-PEG is sufficiently hydrophobic and strongly anchored in the surface bilayer of the LNP to avoid changes in partitioning after protein conjugation independent of the media ionic strength.

The thermodynamics of anchor-PEG-mAb retention or desorption can be modeled by considering the balance between the anchoring energy from hydrophobic interactions that tends toward keeping the molecule attached to an LNP surface, the translational entropy gained from desorption from the surface, and steric penalties that occur from attaching a mAb to an anchor at the LNP surface. We will use *ΔG* > 0 to refer to energies that tend toward retention, and *ΔG* < 0 for energies that promote desorption.

The anchoring energy for PEG-lipids can be estimated from their critical micelle concentrations, *ΔG*_*anchoring*_ = *kTln*(*CMC*), where the CMC is a mole fraction. Literature reports CMC values for DSPE-PEG_2kDa_ of ∼20 uM in DI water and ∼0.5 uM in HEPES-buffered saline, resulting in estimated anchoring energies of 14.8 kT in water and 18.5 kT in buffer.^26^ We use these values to estimate a Δ*g*_*CH*_2__ for the methylene group contribution, Δ*G*_*anchoring*_ = *n*_*CH*_2__ Δ*g*_*CH*_2__, to apply to the PCL_5kDa_-b-PEG_2kDa_, for which CMC values are less readily available. Using *n*_*CH*_2__ = 36 and Δ*G*_*anchoring*_∼ 18.5 kT from DSPE, we estimate Δ*g*_*CH*_2__ ∼ 0.51 kT, which is comparable to values reported elsewhere.^27^ From this we calculate an anchoring energy for the PCL_5kDa_ block (monomer MW 114 Da; 44 monomers, each with 5 CH_2_) around 113 kT, and an energy around 14 kT for DMG-PEG_2kDa_, which is in line with literature values. Nonlinearity in the methylene group contribution term for a large polymer block may require the value for PCL to be adjusted downward, but even a 70% reduction does not affect the following analysis. The PCL block has favorable solubility with the lipid components making up the LNP bilayer, which suggests that the PCL is solubilized into the bilayer rather than collapsed on the surface, supporting the applicability of the methylene group contribution approach.

For translational entropy, we use the ratio of volumes in the bound vs. desorbed states, 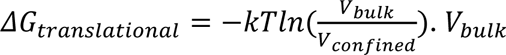 is the molecular accessible volume, 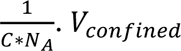 is the molecular volume at the LNP surface; this is estimated as either 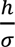 for PEG without a mAb, or 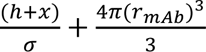 if a mAb is present. *σ* is the PEG surface density, reported in the literature as around 0.273 chains/nm^2^.^5^ *h* is the brush height calculated using de Gennes scaling^28^, *h* = *Na*^5/3^*σ*^1/3^, using number of monomers *N* = 2000Da / (44 Da / PEG monomer) ∼45 monomers, and PEG monomer length *a* ∼ 0.28 nm from literature^29^ ; *h* ∼ 3.5nm. *x* is a small (∼1-2 nm) conformational extension of the PEG tether we allow to ensure mAbs sit above the PEG brush surface, which incurs an energy penalty around -1 kT.^28,30^ Using concentrations *C* for components following FNP, we calculate *ΔG*_*translational*_ ∼-6.74kT for PCL-PEG-mAb and DSPE-PEG-mAb, and *ΔG*_*translational*_ ∼-9.15kT for both unconjugated DMG-PEG or DSPE-PEG.

The steric mAb energy penalty is the sum of three terms. mAbs cannot enter the PEG surface brush, so when tethered at its surface, their rotational conformations are reduced by a factor of 2, since they cannot rotate toward the plane of the brush; this incurs a −*kTln*(0.5)= -0.69 kT penalty. Likewise, the PEG chain end must remain above the brush surface instead of below, a halving of available equivalent microstates that incurs another -0.69 kT penalty. We also include a term for packing spheres on a surface at high volume fraction, *ΔG*_*packing*_ = −*kTln*(1 − *ϕ*) for surface coverage *ϕ*. Since there are roughly 3.5 times as many reactive PEG chain ends on the particle surface (∼830) as the maximum number of 11 nm antibodies that could fit on the surface of a 70 nm diameter LNP (∼233), and due to the size increase by DLS that indicates a very high degree of mAb surface presentation, we estimate this value to be ∼-4.7 kT. Including the -1kT from the small PEG stretching term above, these terms sum to ∼-7.1 kT in favor of desorption.

**Table 2:** Estimated energies contributing to anchor-conjugate retention or desorption.

|  | Anchoring | Steric | Translational | Total | Experimental |  |
| --- | --- | --- | --- | --- | --- | --- |
|  |  |  |  |  | retained<br>*in vitro*? | retained<br>*in vivo*? |
| **DPSE-PEG<sub>2k</sub>-mAb**<br>**(150mM):** | 18.5 kT | -7.10 kT | -6.74 kT | 4.7 kT | ✓ | ✗ |
| **DSPE-PEG<sub>2k</sub>-mAb**<br>**(water):** | 14.8 kT | -7.10 kT | -6.74 kT | 1.0 kT | ✗ | N/A |
| **PCL<sub>5k</sub>-PEG<sub>2k</sub>-mAb:** | 112.7 kT | -7.10 kT | -6.74 kT | 98.9 kT | ✓ | ✓ |
| **DPSE-PEG<sub>2k</sub>:**<br>**(applies also to**<br>**DSG-PEG<sub>2k</sub>)** | 18.5 kT | N/A | -9.15 kT | 9.4 kT | ✓ | ✓ |
| **DMG-PEG<sub>2k</sub>:** | 14.0 kT | N/A | -9.15 kT | 4.8 kT | ✓ | ✗ |

Remarkably, this model recapitulates all reported experimental phenomena observed for retention or shedding of surface anchor-PEG molecules with and without mAb conjugation. It captures the effect of ionic strength on DSPE-PEG_2kDa_-mAb retention during *in vitro* processing, in that the low ionic strength case exhibits a smaller residual anchoring energy (∼1 kT) than may be expected to be required for retention (∼3-7 kT), but at 150 mM ionic strength, the residual ∼4.7 kT for DSPE-PEG_2kDa_-mAb provides sufficient anchoring *in vitro.* This ∼4.7 kT is insufficient to remain anchored *in vivo*, however, and DSPE-PEG_2kDa_-mAb becomes fugitive upon injection – exactly analogous to the ∼4.8 kT residual seen for DMG-PEG_2kDa_, which is also retained *in vitro* but sheds *in vivo*, as previously reported.^8^ Unconjugated DSPE-PEG_2kDa_ exhibits ∼9.4 kT of residual anchoring energy, which is sufficient to remain attached both *in vitro* and *in vivo.* This supports the idea that dilution into the blood, combined with other driving forces for desorption encountered *in vivo*, such as hydrophobic sinks, can overcome at least ∼4.8 kT, but not ∼9 kT, of residual anchoring energy. The residual anchoring energy of the PCL_5kDa_-b-PEG_2kDa_-mAb is well above this threshold.

In this analysis, we considered the mAb as a nanoparticle that rests just above the PEG brush surface, such that it does not exclude any PEG chains beneath it or incur any osmotic pressure penalty. A drawback of these simplifying assumptions is that this version of the model does not explicitly capture a strong dependence on mAb size. Further analysis, supported by molecular dynamics simulations, is needed to develop a robust relationship between the molecular weight of a conjugated protein (mAb, Fab, etc.) and the anchoring energy required to retain it. Nevertheless, the understanding developed here that there exist three regimes for residual anchoring energies – i.e., (1) low energies, wherein the anchor is not retained *in vitro*; (2) intermediate energies, for which the anchor is retained *in vitro* but becomes fugitive *in vivo*; and (3) high energies, where anchors are retained both *in vitro* and *in vivo* – is an important conceptual advancement.

DSPE-PEG_2kDa_-maleimide has shown two unusual behaviors when added to the usual four-component LNP formulation as an antibody conjugation ligand: interference with the fusion-rearrangement of acidic, vesicle-like primary LNPs, and a strong sensitivity of the conjugate stability to the ionic strength of the media. To the best of our knowledge, these behaviors are not common knowledge in the literature on targeted LNPs. When using DSPE-PEGs as part of the LNP formulation, special care must be taken to preserve the same LNP size as conventional, untargeted counterparts and to produce stable, protein-conjugated LNPs. Changes in LNP size during the single-stage dialysis process seem particularly insidious because gross changes in particle diameter can alter the intrinsic biodistribution of nanoparticles *in vivo*, and because larger aggregates could become lost during processing to e.g. filter membranes or dialysis tubing, leading to a false positive size measurement. These adverse behaviors are not unexpected due to the anionic nature of the DSPE-PEG_2kDa_ molecule, where the single negative charge in the head group favors ion pairing with the excess cationic lipids in the formulation and generating negative curvature which destabilizes the LNP surface membrane.^27^

PCL_5kDa_-b-PEG_2kDa_-maleimide block copolymer anchoring of targeting ligands showed improved formulation stability, processability, and *in vivo* stability compared to the DSPE-PEG_2kDa_-maleimide systems. The nonionic, significantly more hydrophobic block copolymer retained mAb and Fab-sized proteins where DSPE-PEG_2kDa_ anchors failed to remain attached to LNP surfaces. These results demonstrate that apparently stable DSPE-PEG_2kDa_ anchored targeting proteins can rapidly dissociate *in vivo* after intravenous administration into the complex, protein-rich blood.

This is a new result that appears to be unexpected in the field of LNP targeting. It is necessary for the field to understand LNP formulation requirements for robust mAb targeting. For example, targeting studies published to date demonstrate targeting to essentially “first pass” targets such as lung epithelial cells^11^ and brain microcapillary epithelial surfaces.^31^ The field of liposomal delivery indicates that circulation times of ∼4 hrs are required for adequate circulation to achieve targeting to lower accessibility targets, such as solid tumors. The short time we observe for the DSPE-PEG_2kDa_-mAbs shedding from LNPs, which is comparable to the rate of DMG-PEG_2kDa_ shedding reported by Cullis, is inadequate to obtain adequate targeting engagement. For the mAbs presented on PCL_5kDa_-PEG_2kDa_-mAb LNPs, the extended circulation time will enable more efficient targeting.

