## Supporting information for "DSPE-PEG does not retain targeting antibodies on LNP surfaces *in vivo*; a higher molecular weight anchor is required"

**Supplemental Information**

**Methods:**

*Antibody conjugation with Holmium tag:*

First, the cetuximab storage buffer was exchanged (30 kDa MWCO centrifugal ultrafilter) for 50 mM bicarbonate at pH 9. Holmium loaded DOTA-Bz-SCN was dissolved in DMSO at 25 mg/mL then added to the cetuximab solution at a molar ratio of 25:1 (Ho)DOTA-Bz-SCN:cetuximab. After three hours the protein solution was again exchanged over a 30 kDa MWCO for 10 mM HEPES at pH 7.4 with 140 mM NaCl. The Ho-tagged protein was then activated for conjugation to the LNP-PEG-maleimide by adding 2 equivalents of TCEP.

*ICP-MS materials and method parameters:*

Materials: Nitric acid trace metal grade (CAS#: A509-P212) was purchased from Fisher Scientific (Waltham, MA, USA). Bovine Liver NIST 1577c, Ethylenediaminetetraacetic acid ACS reagent grade (CAS# 60-004), Hydrogen Peroxide solution (CAS#: 7722-84-1, and Hydrochloric Acid (CAS# 7647-01-0) were purchased from Sigma-Aldrich (St. Louis, MO, USA). Terbium, Europium, Holmium, and Gadolinium standards were purchased from High Purity Standards (North Charleston, SC, USA). 1.5 µm nylon membrane filters (CAS: F2500-12) were purchased from ThermoFisher Scientific (Waltham, MA, USA). ICP-MS tuning solution (CAS: 5185-5959) was purchased from Agilent Technologies, Inc. (Santa Clara, CA, USA)

Methods: ICP-MS analyses were performed using an Agilent 7850 ICP-MS equipped with an x-LENs optics system, MicroMist Nebulizer, PeriPump sample introduction system, and an SPS4 autosampler. Plasma ignition and tuning were conducted in an aqueous Agilent Tune solution. Samples were introduced via the PeriPump at an uptake speed of 0.2 rotations per second (RPS) with an 80-second uptake period, followed by a 50-second stabilization time before data acquisition. After each measurement, the autosampler executed a rinse sequence consisting of a probe rinse for 60 seconds at 0.3 RPS and 30 seconds standard port, and a four-vial rinse cycle, with each vial rinsed for 30 seconds at 0.3 RPS, followed by 30 seconds at the rinse port, ensuring minimal carryover and consistent plasma stabilization across samples. Two tune modes were utilized: No-Gas mode and Helium (He) collision mode. Both modes operated under High Matrix Introduction (HMI) plasma conditions with medium aerosol dilution. No-Gas values were utilized for further analysis due to a slight increase in sensitivity and no concern about spectral interference for the analytes of interest. Additional parameters, such as Plasma Gas conditions, ion optics, and cell parameters, are summarized in **Table S1**. Additional acquisition parameters are listed below in **Table S2**. Both online (Tb) and offline (EU) internal standard recoveries were between 70 and 130%.

Cleaning and chelation procedure for digestion vessels: digestion vessels were first triple-rinsed with ultrapure water. For acid cleaning, 10 mL of 70% trace-metal grade nitric acid was added to each tube. After adding nitric acid, the vessels were sealed and placed in the microwave digestion system, where the Xpress Clean One-Touch program was used to heat and clean the vessels. Following the cleaning cycle, the acid was allowed to cool and discarded, and each vessel was triple-rinsed with ultrapure water. Following the acid wash step a 5% Ethylenediaminetetraacetic acid (EDTA) (w/v) solution was prepared and used as a chelating agent. It is important to note that to dissolve EDTA, the solution was heated to 70 ºC, and 0.1 M NaOH was added dropwise until the suspended EDTA particles dissolved and the solution was clear and particle-free. 10 mL of the EDTA solution was then added to each digestion vessel, where the Xpress Clean One-Touch program was used to heat and clean the vessels. After completion of the cleaning cycle, the EDTA solution was discarded, and the vessels were rinsed in triplicate with ultrapure water. Animal tissue and whole-blood samples were stored at -80ºC before digestion.

**Table S1.** Agilent 7850 ICP-MS instrument operating parameters used for metal quantification.

| **Parameter** | **Value** |
|---|---|
| Radio Frequency (RF) Power | 1600 Watts |
| Radio Frequency Matching | 1.80 Volts |
| Sample Depth | 10 mm |
| Nebulizer (No-Gas) flow | 0.68 $\frac{L}{min}$ |
| Nebulizer (He-Gas) flow | 0.65 $\frac{L}{min}$ |
| Make-up gas | 0.26-0.27 0.68 $\frac{L}{min}$ |
| Spray Champer Temperature | 2ºC |
| Extract 1 | 0.0 V |
| Extract 2 | -200 V |
| Omega lens | 10 V |
| Cell entrance/exit (No Gas) | -30/-50 V |
| Cell entrance/exit (He mode) | -40/-60 V |
| Plate Bias (No Gas) | 14.2 V |
| Plate Bias (He mode) | 1.6 V |
| Octopole bias (No-Gas) | -8 V |
| Octopole bias (He-mode) | -18 V |
| Octopole RF | 200 V |
| Energy discrimination | 5 V |

**Table S2.** The parameters below outline the isotopes, integration times, and detector settings used to quantify lanthanide tracers. Integration times were optimized to balance sensitivity and throughput.

| **Mass** | **Element** | **No-Gas Integration Time (seconds)** | **Helium Mode Integration Time (seconds)** | **Detector Mode** |
|---|---|---|---|---|
| 151 | Europium | 0.6 | 1 | Auto |
| 157 | Gd | 0.6 | 1 | Auto |
| 159 | Terbium | 0.6 | 1 | Auto |
| 165 | Ho | 0.6 | 1 | Auto |

**Results**

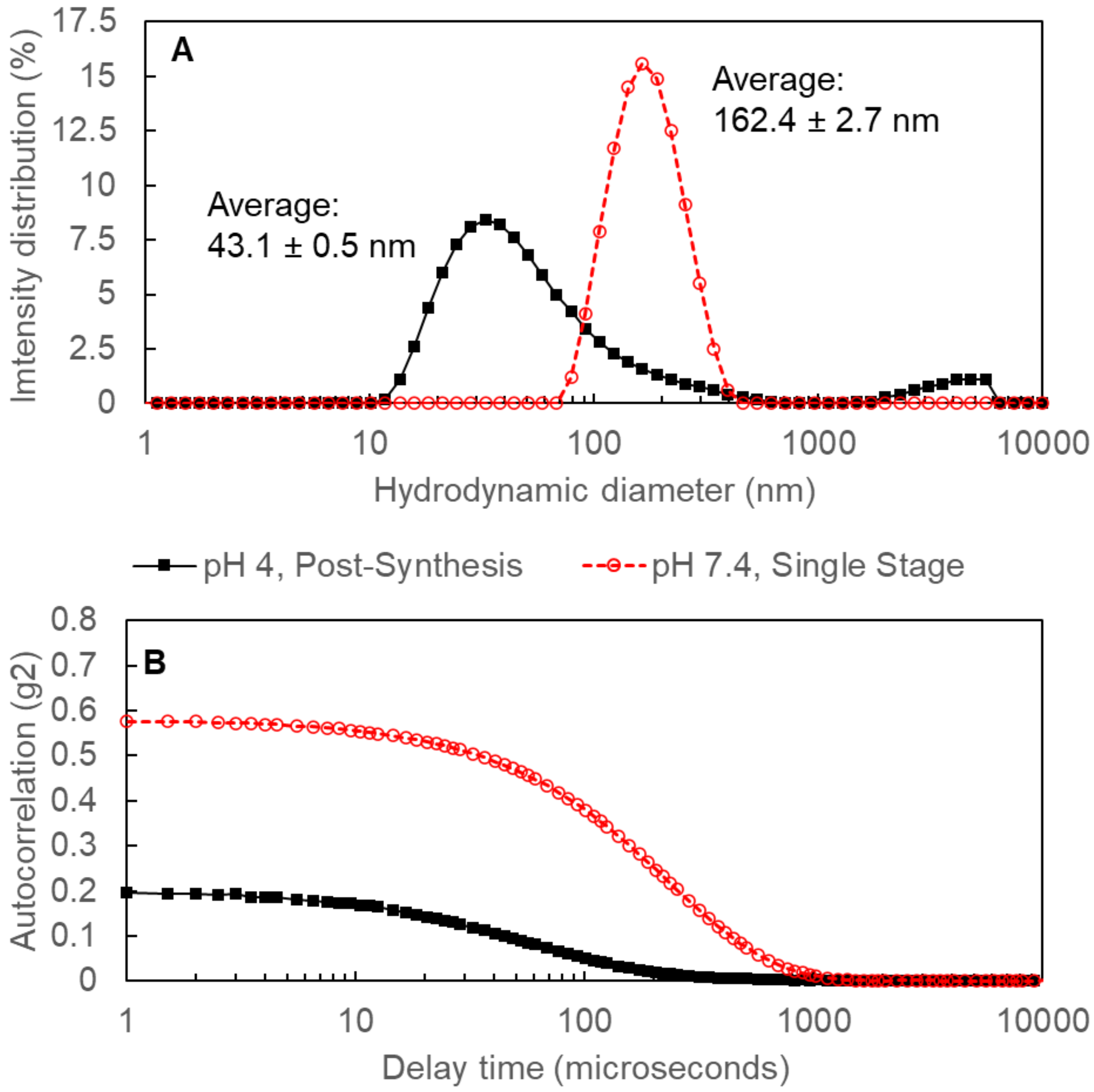


**Figure S1:** When the DSPE-PEG LNPs were directly dialyzed from 10% organic solvent / 90% pH 4 buffer into neutral pH buffer, aggregation to a larger than expected from fusion to recover PEG surface density was observed.

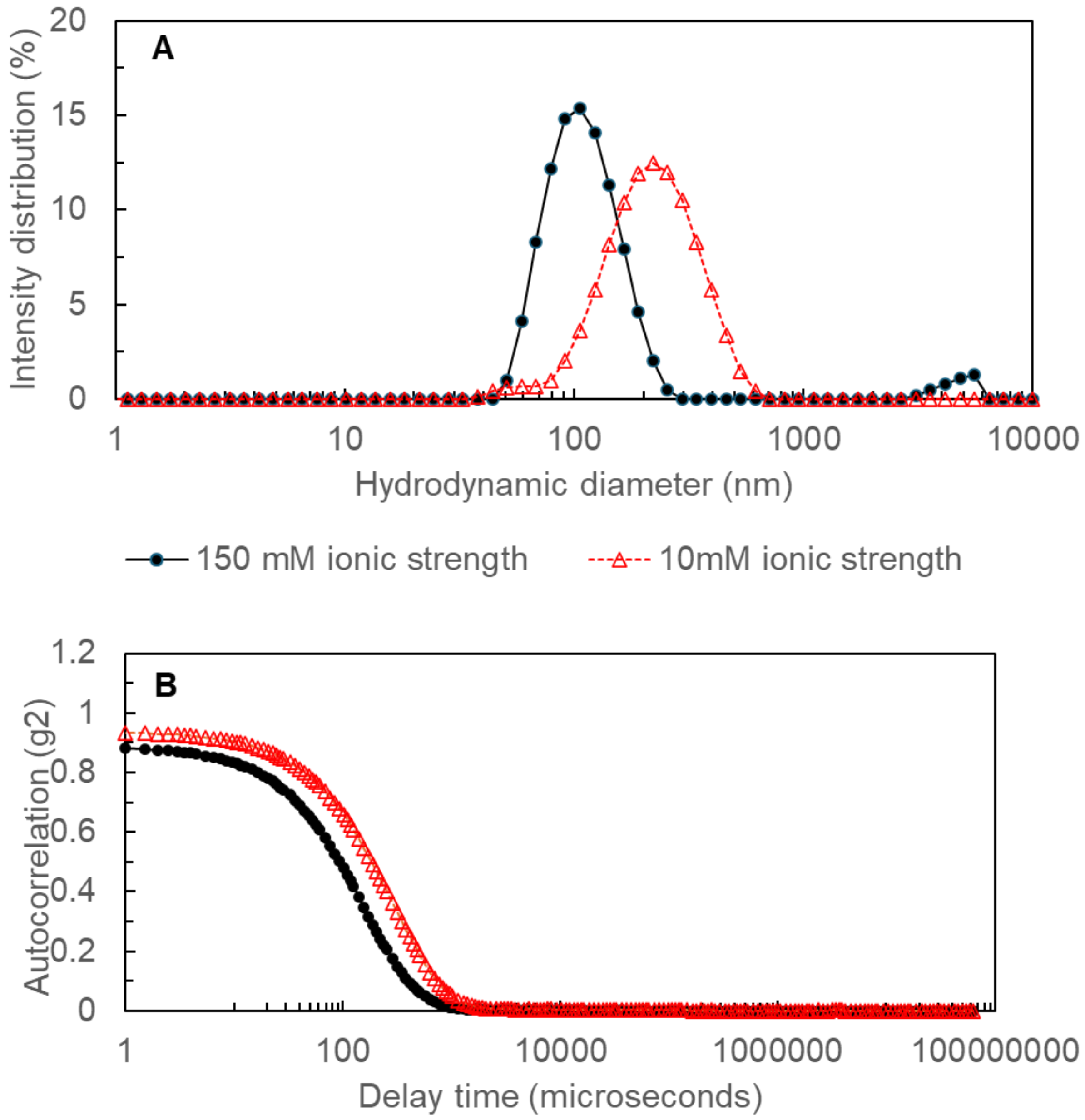


**Figure S2:** When protein conjugation was performed at low ionic strength, a significantly larger increase in LNP size was observed than would be expected (and was observed at high ionic strength) for the addition of a layer of 5-10nm protein to the surface of a 70nm particle.

We note that the strongly hydrophobic PCL block of the PCL-b-PEG can collapse and produce PCL-b-PEG micelles rather than uniformly incorporating PEG into the LNP surface; this would be an expected result if poor (i.e., insufficiently rapid) mixing were used to drive LNP self-assembly. The confined impinging jets mixing used for preparation of all commercial COVID LNPs provides rapid mixing with ~1.5 ms mixing time[16] that enables production of these LNP constructs with PCL-b-PEG. In contrast, laminar flow microfluidic devices may not enable homogeneous incorporation of PCL-b-PEG.[32]

**Table S3**: Measured final organ distributions of Ho and Gd.

| | LNP 1 %ID | | LNP 2 %ID | | LNP 3 %ID | | LNP 4 %ID | |
|---|---|---|---|---|---|---|---|---|
| **Tissue** | **Ho** | **Gd** | **Ho** | **Gd** | **Ho** | **Gd** | **Ho** | **Gd** |
| **Brain, 24h** | 0.01 ± 0.00 | 0.00 ± 0.00 | 0.01 ± 0.00 | 0.01 ± 0.00 | 0.00 ± 0.00 | 0.01 ± 0.00 | 0.00 ± 0.00 | 0.00 ± 0.00 |
| **Heart, 24h** | 0.33 ± 0.02 | 0.06 ± 0.00 | 0.30 ± 0.03 | 0.12 ± 0.02 | 0.00 ± 0.00 | 0.11 ± 0.01 | 0.00 ± 0.00 | 0.08 ± 0.01 |
| **Liver, 24h** | 69.88 ± 18.48 | 133.09 ± 35.88 | 78.88 ± 9.28 | 136.24 ± 16.14 | 0.00 ± 0.00 | 164.87 ± 10.74 | 0.00 ± 0.00 | 172.86 ± 14.64 |
| **Lung, 24h** | 0.33 ± 0.05 | 0.09 ± 0.02 | 0.43 ± 0.13 | 0.25 ± 0.06 | 0.00 ± 0.00 | 0.18 ± 0.03 | 0.00 ± 0.00 | 0.12 ± 0.01 |
| **Spleen, 24h** | 0.80 ± 0.12 | 1.27 ± 0.36 | 0.87 ± 0.19 | 2.38 ± 0.53 | 0.00 ± 0.00 | 1.60 ± 0.28 | 0.00 ± 0.00 | 2.71 ± 1.08 |

Some overage in liver measurements was observed, possibly due to trace metals in rat diets that could not be measured *a priori*.

For a spherical LNP with diameter 70nm, the surface area is $4\pi(35nm)^2 \sim 15{,}400nm^2$. Using the literature value for σ of 0.273 chains/$nm^2$, this surface area corresponds to ~4150 PEG surface chains per LNP, 20% of which, ~830, are terminated in maleimide for protein conjugation.

Spherical proteins pack onto a spherical particle on the surface defined by radius $R_{plane} = R_{particle} + R_{protein}$; we add an additional 1.5 nm to this radius to allow the mAbs to sit fully outside the PEG surface brush layer. Modeling mAbs as 11nm spheres, this $R_{plane}$ = 35 nm + 5.5 nm + 1.5 nm = 42 nm. The area of this spherical plane is $4\pi(42nm)^2 \sim 22{,}100nm^2$. With each mAb occupying a footprint on this surface of $\pi(5.5nm)^2 \sim 95nm^2$/mAb, the maximum number of mAbs that could fit on the surface is $22{,}100nm^2$ / ($95nm^2$/mAb) ~ 233 mAbs. With 830 reactive sites available, the excess is 830/233 ~ 3.5-fold.

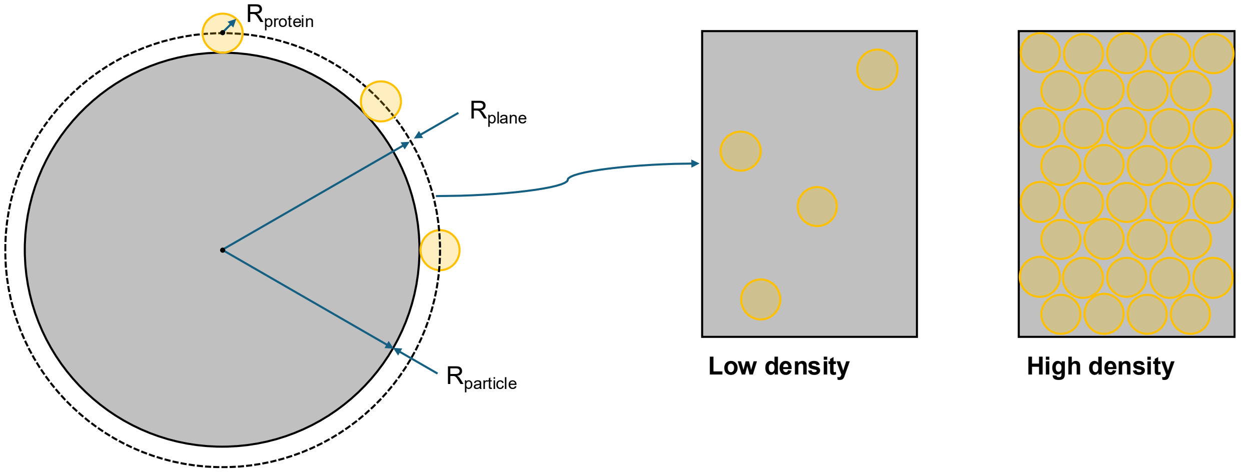


**Figure S4:** Protein conjugation to a nanoparticle surface is fundamentally a particle-particle reaction: proteins are much smaller than the carrier particle. Orange circles are the projected cross-sectional area of the protein.

**Table S4:** Terminal organ time points and the respective ratio of Holmium to Gadolinium for LNP 1 and LNP 2.

| **Tissue** | **(1)** 1.2% DSG-PEG, 0.3% DSPE-PEG-Mal **(Ratio: Gd:Ho)** | **(2)** 1.2% DSG-PEG, 0.3% PCL-b-PEG-Mal **(Ratio: Gd:Ho)** |
|---|---|---|
| Brain | 0.245 $\pm$ 0.048 | 0.644 $\pm$ 0.104 |
| Heart | 0.080 $\pm$ 0.014 | 0.260 $\pm$ 0.019 |
| Liver | 0.951 $\pm$ 0.024 | 1.099 $\pm$ 0.072 |
| Lung | 0.138 $\pm$ 0.025 | 0.374 $\pm$ 0.047 |
| Spleen | 0.785 $\pm$ 0.146 | 1.740 $\pm$ 0.207 |

**Table S5:** Holmium to Gadolinium ratio in each blood sample as a function of time.

| **Formulation** | **1h** | **3h** | **7h** | **10h** | **24 h** |
|---|---|---|---|---|---|
| LNP1 | 0.199 ± 0.019 | 0.135 ± 0.014 | 0.129 ± 0.016 | 0.133 ± 0.024 | 0.158 ± 0.043 |
| LNP2 | 0.649 ± 0.046 | 0.566 ± 0.053 | 0.606 ± 0.102 | 0.606 ± 0.158 | 0.590 ± 0.150 |

**Table S6:** ng/mL of Gadolinium per mL of blood.

| | [Gd] ng/mL blood | | | | |
|---|---|---|---|---|---|
| **Formulation** | **1h** | **3h** | **7h** | **10h** | **24h** |
| LNP 1 | 56.928 ± 17.606 | 9.396 ± 6.533 | 15.720 ± 4.486 | 9.772 ± 2.336 | 4.200 ± 0.788 |
| LNP 2 | 208.420 ± 48.009 | 137.292 ± 25.005 | 75.600 ± 18.103 | 53.784 ± 21.953 | 15.571 ± 4.627 |
| LNP 3 | 243.812 ± 56.845 | 100.120 ± 26.156 | 43.112 ± 6.484 | 27.200 ± 3.173 | 13.882 ± 2.219 |
| LNP 4 | 174.140 ± 30.368 | 66.6 ± 17.413 | 24.784 ± 10.085 | 9.624 ± 3.119 | 2.875 ± 0.271 |

**Table S7:** Gadolinium normalized per gram of tissue.

| | ng Gd / gram tissue | | | |
|---|---|---|---|---|
| **Tissue** | **LNP 1** | **LNP 2** | **LNP 3** | **LNP 4** |
| Brain | 0.194 ± 0.063 | 0.535 ± 0.085 | 0.785 ± 0.160 | 0.224 ± 0.039 |
| Heart | 4.830 ± 1.462 | 14.938 ± 1.699 | 12.432 ± 2.279 | 7.504 ± 1.577 |
| Liver | 1128.055 ± 311.270 | 1446.785 ± 256.786 | 1793.940 ± 174.897 | 1571.673 ± 185.593 |
| Lung | 6.566 ± 1.888 | 22.363 ± 5.614 | 15.904 ± 2.439 | 9.072 ± 0.619 |
| Spleen | 175.943 ± 66.937 | 402.085 ± 80.469 | 285.656 ± 43.630 | 414.036 ± 126.641 |

**Table S8:** Holmium per gram of tissue.

| | Ng Ho metal/ geam tissue | | | |
|---|---|---|---|---|
| **Tissue** | **LNP 1** | **LNP 2** | **LNP 3** | **LNP 4** |
| Brain | 0.794 ± 0.223 | 0.830 ± 0.023 | 0.019 ± 0.006 | <LLOQ |
| Heart | 59.365 ± 8.860 | 57.633 ± 6.540 | 0.047 ± 0.000 | <LLOQ |
| Liver | 1182.269 ± 314.237 | 1319.486 ± 237.755 | 0.009 ± 0.021 | 0.003 ± 0.007 |
| Lung | 46.900 ± 6.939 | 60.312 ± 15.102 | 0.006 ± 0.012 | <LLOQ |
| Spleen | 217.625 ± 47.360 | 231.812 ± 47.028 | 0.023 ± 0.000 | <LLOQ |
